# Numerical Solution to SIR Model using SBP-SAT operators

**DOI:** 10.1101/2025.09.22.677686

**Authors:** Sam Motsoka Rametse, Sheldon Herbst

## Abstract

Accurate and stable numerical simulation of epidemic dynamics is essential for translating mathematical models into reliable computational tools for public health. The Susceptible–Infectious–Recovered (SIR) model remains a cornerstone of mathematical epidemiology, yet the robustness of its numerical treatment strongly influences predictive reliability in applications ranging from outbreak forecasting to intervention assessment. Here, we introduce a high-order Summation-By-Parts (SBP) framework with Simultaneous Approximation Terms (SAT) for the numerical solution of the SIR system. Unlike standard integrators, the SBP-SAT method enforces stability at the discrete level while retaining high accuracy, offering a principled approach for handling nonlinear epidemic dynamics. We validate the scheme against the analytical solution of the classical SIR model, demonstrating both accuracy and robustness. By situating SBP-SAT methods within the growing landscape of numerical approaches for SIR-type models—including Runge–Kutta, finite difference, and fractional-order solvers—this work establishes SBP-SAT as a powerful and generalizable alternative for computational epidemiology. Beyond the classical SIR model, these results pave the way for applying SBP-SAT schemes to more complex epidemic models where stability and accuracy are critical, thereby advancing the methodological foundations of computational biology.

## 1. Introduction

Mathematical modeling infectious diseases can be traced back to Daniel Bernoulli in the 1760s, when he presented a mathematical model solution to smallpox. The first fundamental mathematical model for epidemic diseases was formulated by Kermack and McKendrick in 1927 [9]. This model applies for epidemics having a relatively short duration (compared to life duration) that take the form of a sudden outbreak of a disease that infects (and possibly kills) a substantial portion of the population in a region before it disappears” [4]. SIR models are based on the mass-action principle introduced by [2], which assume that the course of an epidemic depends on the rate of contact between the susceptible and the infected individuals.There are various developments of the epidemic models to multicompartment models [1, 7, 6, 11].

While the theoretical foundations of SIR models are well established, their practical utility increasingly depends on numerical simulation. Classical methods—such as Runge–Kutta schemes, finite difference approaches, and fractional-order solvers—have been widely applied, but challenges remain in balancing accuracy, stability, and efficiency, particularly for stiff or nonlinear epidemic dynamics. These limitations can affect both the reliability of forecasts and the interpretability of model outcomes in computational biology.

In this study, we address these challenges by introducing a high-order Summation-By-Parts (SBP) framework with Simultaneous Approximation Terms (SAT) for the numerical solution of SIR models. This approach enforces discrete stability while retaining accuracy, providing a principled alternative to conventional schemes. By situating SBP-SAT methods within the broader landscape of numerical epidemic modeling, we aim to advance the methodological toolkit available for computational epidemiology.

In this model, the population is divided into three groups:

- Individuals who are not infected but susceptible (S) to the disease
- Individuals who are already infected(I) with the disease
- Individuals have recovered (R) or removed (dead) from the disease

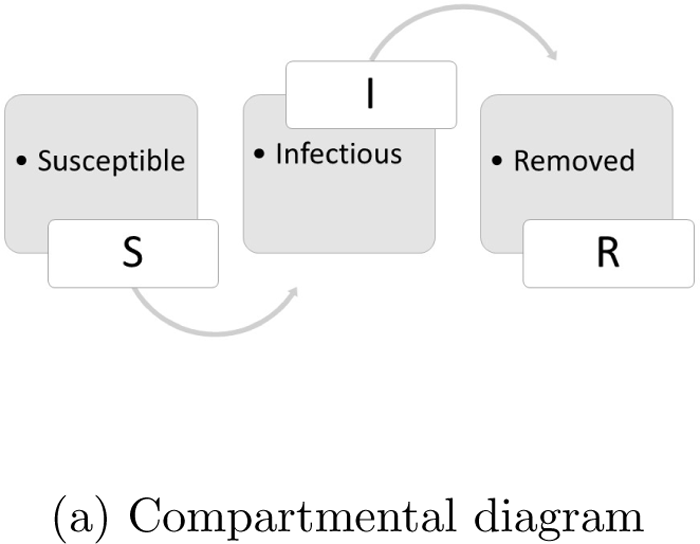

The total population is

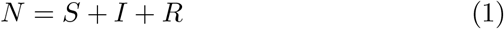

We assume that the population does not change with time

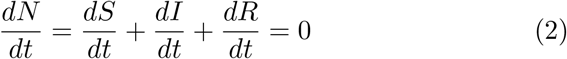

Let *κ* be the actual number of individuals a susceptible person interacts with and *ρ* be the probability that a susceptible gets infected at contact with an infected individual. The rate at which the susceptible group gets reduced with time is given by

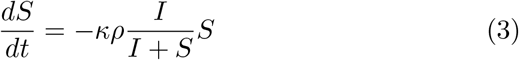

The infected group increases at same rate, but also some individuals get removed from this group.

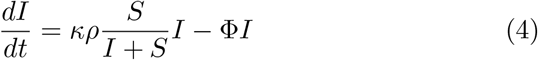

The rate at which individuals get removed (recover or die) from the total population is given by

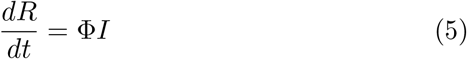

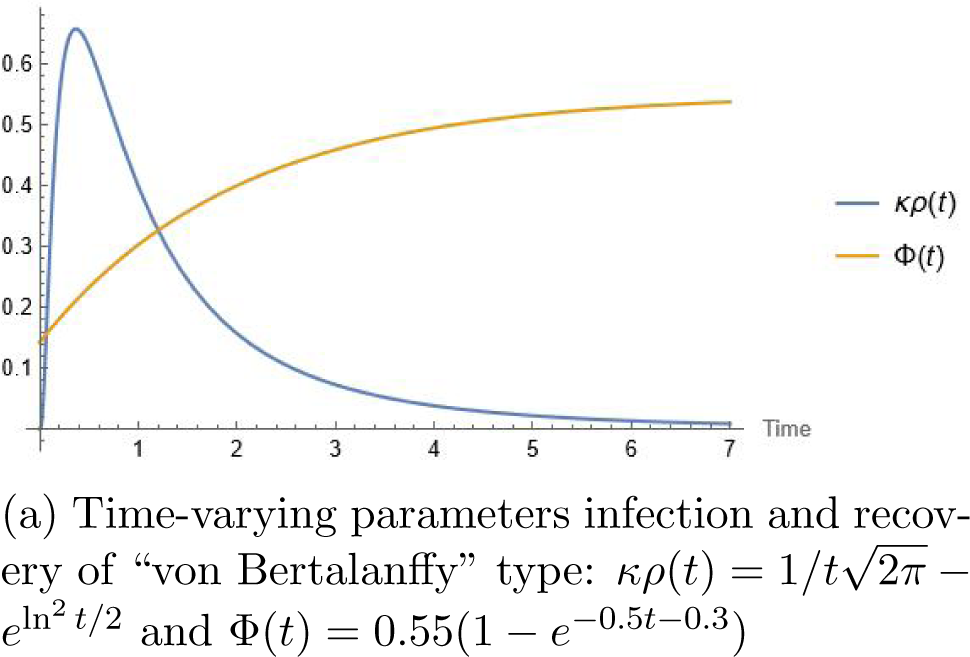

### 1.1 Continuous SIR Model

The model is a system of three differential equations given by

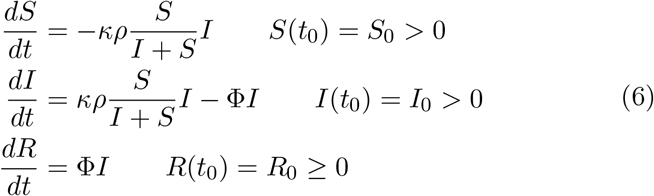

We can derive the exact solution to this system of equations based on the work of Bohner [3]. The first two equations can be re-written as

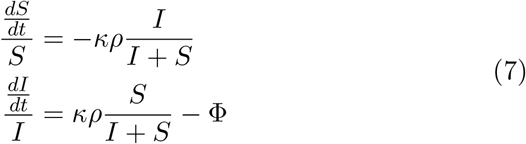

Allowing *β* = *κρ* and combining the two differential in (6) we get

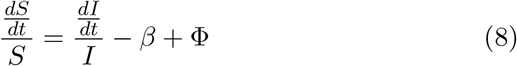

Which is equivalent to

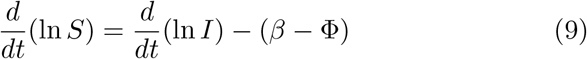

Integrating both sides from 0 to *t* we get

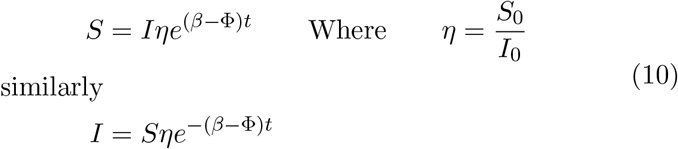

If *β* ≠ Φ, then substituting in (6) we get a differential equation

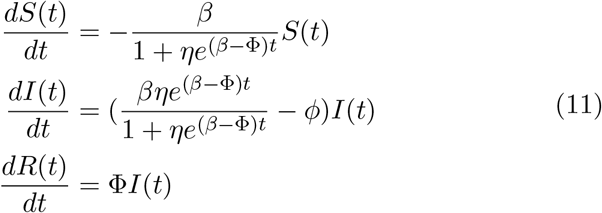

The first order differential equation of (11) can be written as

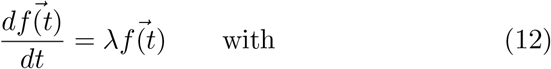

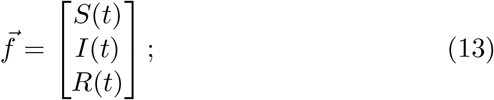

and

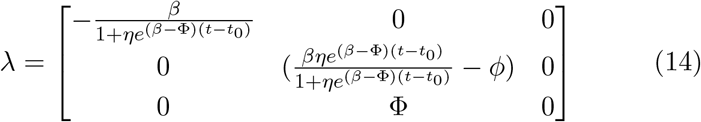

with initial conditions 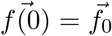 and 0 ≤ *t* ≤ *T*.

## 2 Numerical formulation of the SIR Model

The numerical solution of the SIR model, a system of ordinary differential equations, is crucial for simulating and predicting disease spread.Due to the model’s nonlinearity, analytical solutions are often unattainable, necessitating numerical methods. Various techniques are employed, including the Euler method [14] and Runge-Kutta methods[12]. Below we propose a novel approach of using the SBP-SAT method to solve SIR model.

The Summation by Parts (SBP) property [10, 13, 8] for the first-derivative difference operators, and the Simmulteneous Approximation Term (SAT) technique for weakly imposing boundary conditions[5]. The SBP-SAT method is a powerful yet simple method that allows you (by construction) to exactly mimic the underlying continuous energy estimate in a semi-discrete setting, thus proving stability, which in turn imply convergence.

### Definition 2.1

*We define D an n-dimensional square matrix as the (SBP) finite difference operator for the first derivative as* 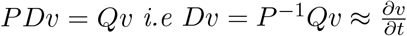

1. *Dt*^*k*^ = *kt*^*k*−1^.
2. *P* = *P* ^*T*^ > 0, *which implies that P is positive definite*.
3. *D* = *P*^−1^*Q*
4. *D*1 = 0, *where both* 1 *and* 0 *are (nx1) vectors*
5. *Q* + *Q*^*T*^ = *E*_*N*_ − *E*_0_

Q is an almost skew-symmetric matrix allowing for the following expression

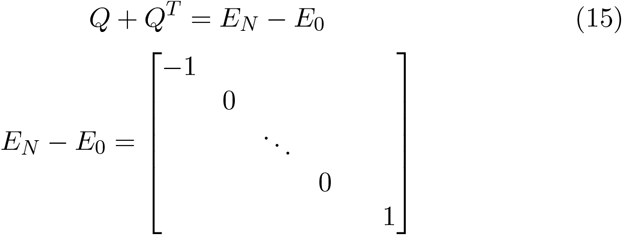

The matrices *D, P* and *Q* are given by

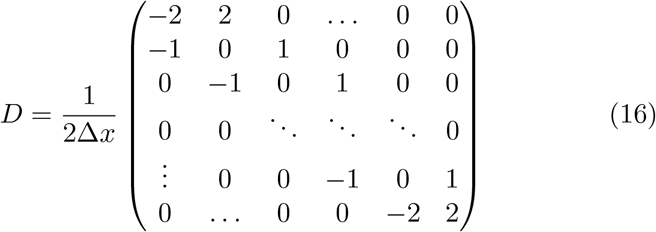

and

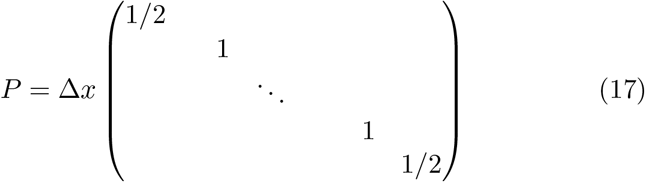

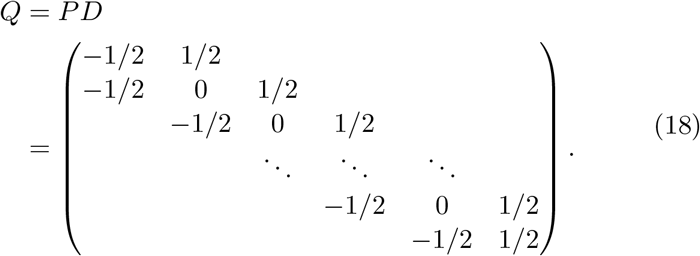

Because P is positive definite we can define the following *norm* and *inner product*

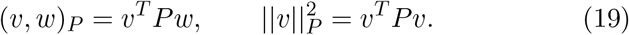

Based on (19) we can show that the discrete analog of integration by parts, (*u, v*_*t*_) = *uv* − (*u*_*t*_, *v*), holds for SBP operators.

### Proposition 2.1

*If D is an SBP operator as defined above, then* (*u, Dv*)_*P*_ = *v*_*n*_*u*_*n*_ − *v*_0_*u*_0_ − (*Dv, u*)_*P*_

*Proof*.

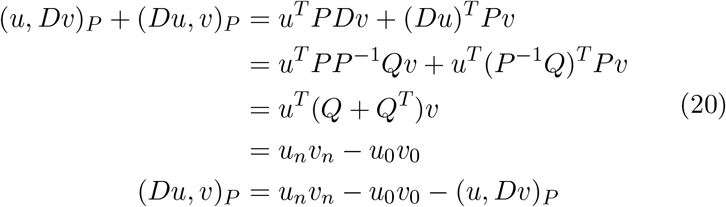

### 2.1 Simultaneous Approximation Terms

To be able to approximate the solutions numerically and still obey boundary conditions we introduce Simultaneous Approximation Terms (SATs). The basic idea behind the SAT technique is to impose the boundary conditions weakly, such that the SBP property is preserved and an energy estimate can be obtained.The SAT acts as a penalty pulling the boundary condition toward the initial conditions. When the boundary conditions are obeyed, the SAT has no contribution. We now impose boundary conditions through the introduction of *S*, the SAT terms given by *S* = *e*_*L*_(*S*_0_, 0, 0, …, 0), with *e*_*L*_ = [1, 0, 0, …, 0]^*T*^ and *S*_0_ = *σ*(*v*_0_ − *f*_0_), *σ* is a parameter that we tune for stability as will be demonstrated below. The discretised version of (11) can be written as

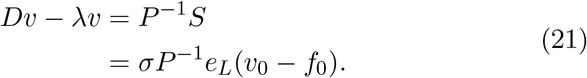

The penalty term in 21 forces the discrete solution towards the initial data. *v*_0_ ≈*f*_0_, but never equal. We use this method to preserve the SBP property of the difference operator *D* = *P*^−1^*Q* is important for the stability proof.

A common way of demonstrating well-possedness of equation (21)is through the *energy method* through the following steps.

1. Multiply the given pde by the solution.
2. Integrate in space.
3. Manupulate the remaining integrals using integration by parts
4. Apply boundary conditions
5. Integrate in time

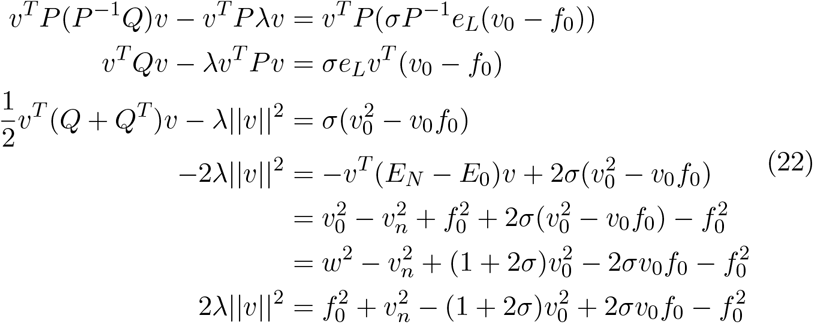

The choice *σ* ≤ −1*/*2 makes the discretization consistent.

## 3 Numerical examples

Let us consider equation 21 can be written as

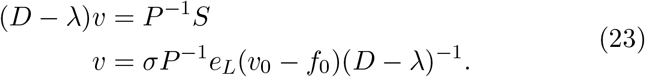

We begin by dividing the temporal 0 ≤ *t* ≤ *T* into *n* + 1 equally distanced grid points, {*t*_*j*_ = *j*Δ*t, j* = 0, 1, 2,.., *n*, Δ*tn* = *T*} where Δ*t* = 1*/n*. We define the vector **v**(*t*) = [*v*_0_, *v*_1_, …, *v*_*n*_]^*T*^ as an approximation of the continuous solution *u*(*t*) at each point *t*_*j*_.

Taking *t* = 3 for the susceptible population 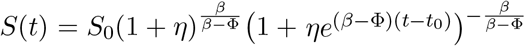 and comparing with the SBP-SAT numerical solution, we see that we achieve order 2 conversion rate.

| Error Analysis with S ≈ 0.4999 | | | |
| --- | --- | --- | --- |
| n | Approximation | Error | Conversion rate |
| 15 | 0.399006 | 0.100984 | 2.46431 |
| 30 | 0.397742 | 0.102248 | 2.167757 |
| 60 | 0.39114 | 0.102876 | 1.970063 |
| 120 | 0.3968 | 0.103195 | 1.830949 |
| 240 | 0.396644 | 0.103346 |  |
| Error Analysis with I ≈ 0.4 | | | |
| n | Approximation | Error | Conversion rate |
| 15 | 0.383561 | 0.0164394 | 2.464018 |
| 30 | 0.382296 | 0.017704 | 2.167289 |
| 60 | 0.381667 | 0.018333 | 1.970063 |
| 120 | 0.381353 | 0.018647 | 1.82915 |
| 240 | 0.381196 | 0.018804 |  |
| Error Analysis with R ≈ 0.35 | | | |
| n | Approximation | Error | Conversion rate |
| 15 | 0.388778 | 0.038778 | 2.454224 |
| 30 | 0.387479 | 0.037479 | 2.159449 |
| 60 | 0.386833 | 0.036833 | 1.963918 |
| 120 | 0.386511 | 0.036511 | 1.82436 |
| 240 | 0.38635 | 0.03635 |  |

| Conversion Rates Comparison using $S \approx 0.4999$ | | | |
| --- | --- | --- | --- |
| n | Euler | Runge-Kutta 4 | SBP-SAT |
| 15 | 1.22113 | 1.215450 | 2.46431 |
| 30 | 1.03144 | 1.026532 | 2.167757 |
| 60 | 0.910035 | 0.90579 | 1.970063 |
| 120 | 0.824026803 | 0.8061 | 1.830949 |

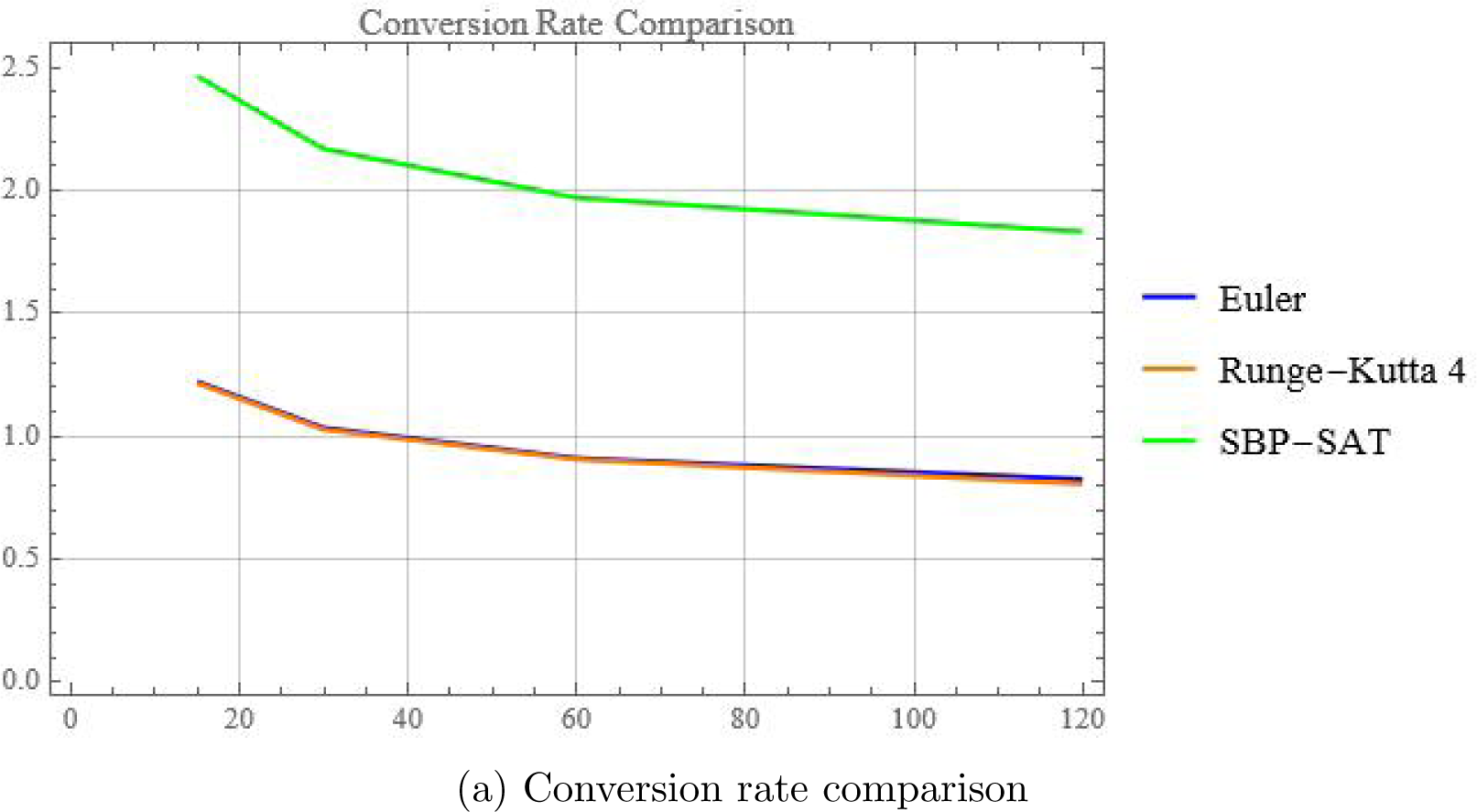

## 4 Discussion

The stable Summation by Parts (SBP) scheme, with a convergence rate of two, has demonstrated its effectiveness in numerically solving the classical SIR epidemiological model. This method, which incorporates Simultaneous Approximation Terms (SATs) to enforce boundary conditions, offers significant advantages for the study of disease dynamics.

A key benefit of the SBP-SAT scheme is its guaranteed energy stability. The SBP framework, when used with boundary conditions enforced by SATs, prevents non-physical oscillations and ensures that the numerical solution remains bounded and physically realistic. This is a critical feature for epidemiological models, where maintaining the physical constraints of the system is essential to prevent misleading conclusions. The stability of the method ensures that the numerical solution accurately reflects the true behavior of the SIR model, providing reliable insights into disease progression.

## 5 Conclusion

In this paper, we have successfully implemented a stable Summation by Parts (SBP) scheme with simultaneous approximation terms (SATs) and a second-order convergence rate to solve the SIR epidemiological model. The numerical results demonstrate the method’s accuracy, stability, and efficiency in capturing the essential dynamics of disease spread. The stable and accurate numerical solution provided by this method contributes to a deeper understanding of epidemiological phenomena and can aid in the development of effective public health strategies.

## Notes

### Competing Interest Statement

The authors have declared no competing interest.

